# *In silico* and *in vitro* characterization of the mycobacterial protein Ku to unravel its role in non-homologous end-joining DNA repair

**DOI:** 10.1101/2023.06.07.543977

**Authors:** Joydeep Baral, Gourab Bhattacharje, Sagarika Dash, Dibyendu Samanta, Elizabeth Hinde, Isabelle Rouiller, Amit Kumar Das

**Author notes:** Corresponding authors: (A.K.Das) (I.Rouiller) (E.Hinde). Authors contributed equally (Gourab Bhattacharje, Sagarika Dash).

## Abstract

Non-homologous end-joining DNA repair is essential for the survival and sustenance of *M. tuberculosis* (Mtb) in the dormant stage of its life cycle. The ability of Mtb to sustain itself in the inactive form has been reported to be the critical factor for its resilience over the years. To unravel one of the salient features of the Mtb’s arsenal, we exploited *in silico* and *in vitro* tools to characterize the DNA binding properties of mycobacterial protein Ku (mKu) and its role in mycobacterial NHEJ. Here, we report the strong affinity of mKu for linear dsDNA exhibiting positive cooperativity for dsDNAs (≥40bp). Molecular dynamics complemented with *in vitro* experiments showed that the DNA binding of mKu provides stability to both mKu homodimer and the DNA. Furthermore, mKu end-capping of DNA was seen to protect the DNA termini against nucleolytic degradation by exonuclease. The DNA-mKu association formed higher-order oligomers probably due to the lodgement of two DNA molecules at opposite ends of the mKu homodimer. The ability of mKu to form continuous filament-like structures with DNA indicated its potential role in mycobacterial NHEJ synapsis.

## 1. Introduction

DNA damage is an inevitable phenomenon for all living systems. Among the various forms of DNA damage, Double-Stranded DNA Break (DSB) plays a pivotal role in cellular life, owing to the detrimental ability of a single unrepaired DSB to result in cell death [1]. Unlike single-stranded DNA breaks, the absence of an undamaged template makes DSBs critical and its repair error-prone. Further, the physical detachment of the broken DNA strands makes it challenging for the cellular DNA repair machinery to re-establish the original template. DNA double-stranded breaks are readily caused by both endogenous (free radicals and reactive species) and exogenous factors (Ionizing radiation, UV-A, and synthetic chemical agents) [2]. Thus, to combat the DSBs and avoid disruption of cellular homeostasis, living systems employ two main DNA repair mechanisms-homologous recombination(HR) and non-homologous end joining(NHEJ) [3]. In response to DNA damage, both repair pathways have been reported to be inter-complimentary in the absence of the other [4]. Even though HR and NHEJ contribute almost equally, their mode of action differs in many vital aspects.

Compared to NHEJ, HR is less error-prone and can efficiently keep the integrity of the genome intact. Conversely, during cellular dormancy or the non-replicating phase of the cell cycle, the homology-dependent repair mechanism comes to a halt due to the absence of a homologous DNA template. In this circumstance, the NHEJ pathway is considered the only alternate damage response being a template-independent repair process. NHEJ repair machinery is considerably fast and versatile throughout the cell cycle but has a trade-off of INDEL (insertion-deletion) mutations in the genome. Indel mutations are often alarming but crucial for genomic diversity and natural selection [5]. For the same reason, apart from DSBR, essential cellular processes such as V(D)J recombination rely on classical NHEJ for the evolution of antibodies [6].

In line with the eukaryotes, many prokaryotic species possess a conserved NHEJ apparatus to repair DSBs [7]. Relevance and importance of NHEJ across microbial kingdoms though similar but are more profound in pathogenic forms. Since a large section of pathogenic and spore-forming microbes spend a significant portion of their life cycle in dormancy (non-replicating phase), thus NHEJ works as an indispensable DSB repair tool affecting their pathogenesis and sustenance. Intracellular pathogens like *Mycobacterium tuberculosis* employs its ability to sustain in an inactive state within the host alveolar macrophages. This population of dormant *M.tuberculosis* residing within macrophages may later transforms into resistant persister cells [8, 9]. For the long-term tenancy in a genotoxic environment constantly threatened by the host’s noxious ROS/RNS species, a fast, template-independent, low-fidelity DNA repair system is obligatory [9]. Though both eukaryotic and prokaryotic NHEJ machinery shares many common features, the latter is less complex regarding the players involved in the process. The comparative simplicity is solely the result of multifunctional enzymes capable of executing the otherwise multi-protein repair process.

The mycobacterial NHEJ apparatus comprises of two key members: Mycobacterial Ku (mKu) and Ligase D (LigD). In the account of a DSB event, mKu serves as the DNA end binding protein, interacting with blunt and staggered-ended DNA with similar affinity but not to closed circular or single-stranded DNA[10]. As bacterial genomic DNA is circular and linear DNA can only be produced by DSB. Thus, the selective affinity of mKu for linear over the circular form of DNA can be speculated to act as a DSBR switch, substituting for the lack of a sophisticated DSB signalling cascade, as seen in eukaryotes. Additionally, the capping of DNA ends by mKu instantaneously after DSB, may protect the exposed DNA termini, which are otherwise damage prone. Similarly, the human form of Ku (Ku70/80) has been reported to preserve 3’- and 5’-terminal overhang regions of double-stranded DNA from DNase I digestion *in vitro* [11]. The second member of the pathway, ligase D, is a multifunctional enzyme harbouring a unique variety of nucleotidyl transferase activities as primase, polymerase, terminal transferase, and 3′ to 5′ exonuclease [12]. Mycobacterial sp. also possesses other ligases, such as, ligase A (NAD+ dependant), ligase B and C (ATP dependant), which are found to associate with mKu [13]. Unlike LigD, these enzymes lack multifunctional domains and have marginal activity in the absence of LigD. Thus mKu is considered the sole activator of mycobacterial NHEJ repair, with LigD as its predominant partner [9]. Among the two key players, ligase D has been characterized in great detail due to the availability of its crystal structure. Owing to the lack of structural information on any prokaryotic Ku, its DNA binding properties are yet to be unravelled.

Across species harbouring an active NHEJ, Ku is found to have conserved structural and functional features. The reason behind this may be referred to the origin and evolution of the protein from species to species. The evolutionary origin of mKu can be traced back to mycobacteriophages (Omega and Corndog phage) [14]. The amino acid sequences of the phage and mycobacterial Ku homologues are highly similar, suggesting probable lateral gene transfer events [14]. The similarity is even consistent with its homologue Gam, which may have transcended from bacteriophage Mu to *E.coli*. Thus, Ku is claimed to have originated from viruses/bacteriophages, which by subsequent gene transfer events, gained additional abilities suited to the requirements of the harbouring species [14]. In Omega and Corndog phages, which lack an active NHEJ machinery, the role of Ku is focused not only on DNA stabilization but also phage genome circularization [15]. To overcome the lack of an active NHEJ, these phages are reported to harness the repair machinery of the host (*Mycobacteria* sp.) complimenting with its own Ku [14, 15] for circularization and later packing of its genome. Throughout evolution, the primordial homodimeric form of the protein Ku has been subjected to the loss and gain of additional domains [14]. Besides the modifications of Ku through evolution from its primordial form in viruses to eukaryotes, the core DNA binding region (cDBR) of the protein has remained conserved. Structure based phylogenetic analysis on Ku has revealed the presence of cDBR across all forms with identical structural conformations, owing to the DNA binding property of Ku [16], wherein, the removal of the cDBR results in comparative loss of its DNA binding property.

In addition to the conserved cDBR, the C-terminal tail of Ku also shows sequence non-specific DNA binding affinity. However, there is no report of any sequence or conformation-based consensus across species. In the case of Mycobacteria, the C-terminal tail has been found to possess tandem PAKKA amino acid repeats responsible for DNA binding [17]. However, Ku from pathogenic mycobacterial strains such as *M.tubeculosis*, *M.bovis*, etc., lack this tandem signature repeat, thereby creating a knowledge gap corresponding to its DNA binding properties. Thus this study, delineates the molecular behaviour of prokaryotic Ku toward its DNA substrate using *in silico* and *in vitro* tools.

## 2. Materials and methods

### 3.1. Sequence analysis

Nucleotide and amino acid sequences of mKu (Rv0937c) were obtained from UniProtKB database [18]. The physico-chemical properties were analysed using Expasy proteomics server [19]. Multiple sequence alignments were performed to determine the conserved sequences among protein homologues using Clustal omega server [20]. Three dimensional structure files were obtained from RCSB PDB database [21].

### 3.2. Molecular modelling

Homology modelling was performed in Swiss-Model [19, 22] and Phyre2 [23] web servers. Ab-initio modelling was performed in Robetta [24] and I-TASSER [25] web servers. The generated models were validated by Ramachandran plot analysis, using Molprobity web server [26]. Comparative analysis among predicted models was performed considering several parameters, such as-average root mean square deviation (RMSD) with conserved Ku70 core DNA binding domain (PDB-1JEY) [27], total potential energy and Z-score, to assess the overall model quality (S1 Table 1). Superimposition of conserved DNA binding core domain and its respective RMSD calculation was performed using Chimera [28]. Total potential energy was calculated using energy minimization of Gromacs [29] and Prosa-web server [30] was used to determine the Z score of the respective models.

### 3.3 Model refinement and protein-protein docking

Water based model refinement was performed using HADDOCK Refinement interface [31]. The refined models were further energy minimized using Gromacs software for molecular docking. The energy minimized mKu models were submitted to multiple servers for molecular docking, HADDOCK [32], Cluspro [32] and ZDock [33] for comparative analysis (S1 Table 2). MMGBSA (Molecular mechanics with generalized Born and surface area solvation) calculations of the docked structures were performed on HawkDock [34, 35] web server. The docked structures were screened on the basis of lowest binding free energy obtained from MMGBSA analysis.

### 3.4 DNA-protein docking

Three different lengths of B-form DNA (20bp, 29bp and 40bp) were used for mKu homodimer-DNA docking (S1 Table 3). The GC content of the DNA molecules were kept at 65.5%, to mimic the mycobacterial genomic DNA. Considering the sequence non-specificity of mKu, the nucleotide sequence of DNA substrates were generated using random sequence generator. All energy minimisations were performed in Gromacs. Minimised models for mKu complex and DNA molecules were docked at HDOCK web server [36]. Docked structures were screened and selected based on their respective docking score (HDOCK) and potential energies (Gromacs).

### 3.5 Molecular Dynamics (MD) simulation

Molecular dynamics (MD) simulations of (mKu)_2_-DNA_(65.5%GC)_ complex were carried out using Gromacs on Ubuntu 18.04 Bionic Beaver. Simulations were performed for 100ns and the trajectory data was collected at every 10ps. AMBER99SB protein and AMBER94 nucleic acid forcefields [37], and TIP3P water model were used for all the simulations. A rectangular box was used as a unit cell for periodic boundary condition. Packmol package [38] was used to calculate the dimension of the rectangular box and the number of ions required to maintain a neutralized solution with physiological NaCl concentration of 0.16M. Steepest descent algorithm was used for energy minimizations with maximum force F_max_ not exceeding 1000 kJ/mol/nm. Each system was equilibrated at temperature 300K and pressure 1 bar by two consecutive 100 ps simulations with canonical NVT ensembles and isobaric NPT ensembles respectively. Protein mKu and the docked DNAs were coupled together for position restraint and thermostat coupling. Trajectories generated from the MD simulations were analysed using Gromacs tools. The obtained data was graphically represented using GraphPad Prism version 9.0.

### 3.6 Recombinant protein production

The bacterial expression plasmid coding for mKu (Rv0937c) was received as a kind gift from Prof. Aidan Doherty, University of Sussex. The cloned plasmids were sequenced, isolated and re-transformed into chemically competent BL21 (DE3) C43 *E.coli* cells. The resultant clones were grown in Luria Broth supplemented with 100 µg.ml^-1^ ampicillin at 37 °C until OD_600_ reached 0.6, followed by induction with 0.25 mM IPTG (Isopropyl β-D-1-thiogalactopyranoside) for 16 hours at 16 °C for over-expression.

The cells from 4ltr culture were harvested after and resuspended in buffer A (50 mM Tris pH 7.5, 250 mM NaCl, 10% Sucrose, PMSF 17 μg.ml^-1^, Benzamidine 34 μg.ml^-1^) followed by addition of Triton X-100 to 0.1% and lysozyme to 0.1 mg.ml^-1^ and finally kept incubated ice for 20 mins. The suspension was lysed by ultrasonication on ice and the lysate was centrifuged at 14000 rpm for 40 min at 4 °C. The resultant supernatant was treated with PEI (Polyethyleneimine; Sigma) to remove DNA contaminants. 5% (v/v) PEI was added in a dropwise manner under constant stirring at 4 °C till the solution reached final concentration of 0.15% (v/v) PEI. Following PEI precipitation, the turbid solution was centrifuged at 14000 rpm for 40 min at 4 °C to remove the precipitants.

The obtained supernatant was incubated with pre-equilibrated Ni-Sepharose high performance affinity matrix (GE Healthcare Biosciences) in buffer A for 16 hours at 4 °C in a test tube rotor at 5 rpm. The incubated sample was loaded on to an empty column and extensively washed with buffer A followed by buffer B (50 mM Tris pH 7.5, 400 mM NaCl, 10mM imidazole, 10% Glycerol, PMSF 17 μg.ml^-1^, Benzamidine 34 μg.ml^-1^) and buffer C (50 mM Tris pH 7.5, 400 mM NaCl, 45mM imidazole, 10% Glycerol, PMSF 17 μg.ml^-1^, Benzamidine 34 μg.ml^-1^) to remove non-specifically bound contaminants. Finally, the protein of interest was eluted with buffer D (50 mM Tris pH 7.5, 400 mM NaCl, 300mM imidazole, 10% Glycerol, PMSF 17 μg.ml^-1^, Benzamidine 34 μg.ml^-1^). The eluent from IMAC was desalted in buffer E (50 mM Tris pH 7.5, 50 mM NaCl, 2mM DTT 10% Glycerol, PMSF 17 μg.ml^-1^, Benzamidine 34 μg.ml^-1^) for Ion-exchange chromatography (IEX). The anion exchange was performed using Q-Sepharose matrix (GE Healthcare Biosciences) on an ÄKTA prime Plus system (GE Healthcare Biosciences) pre-equilibrated with buffer E. Fractions of 2 ml were collected at a flow rate of 2 ml/min under a continuous gradient of NaCl (50mM to 600mM). The fractions containing the desired protein were pooled together and concentrated to 2ml using a 10 kDa cutoff Vivaspin20 concentrator (GE Healthcare Biosciences). Then the protein was subjected to size-exclusion chromatography using HiLoad Superdex 200 prep-grade 16/600 column (GE Healthcare Biosciences) pre-equilibrated with buffer F (50 mM Tris pH 7.5, 150 mM NaCl, 2mM DTT 10% Glycerol, PMSF 17 μg.ml^-1^, Benzamidine 34 μg.ml^-1^). Fractions of 2 ml were collected at a flow rate of 1 ml/min on an ÄKTA prime Plus system (GE Healthcare Biosciences) equilibrated with buffer F. The fractions containing the desired protein were pooled together. The protein concentration was estimated from the absorbance at 280 nm as well as by the Bradford method [39] and the purity was verified by 12% SDS PAGE.

### 3.7 Differential scanning fluorimetry (DSF)

In order to validate the impact of DNA binding on the thermal stability of the protein mKu we exploited DSF to determine the melting temperature (Tm) of mKu in presence and absence of 40bp dsDNA. For this experiment we have referred to the previously published protocols to determine the Tm of a protein using a solvatochromic dye - Sypro orange (Invitrogen) as a reporter [40, 41]. The dye SYPRO orange has marginal fluorescence in solution but it fluoresces upon binding to the hydrophobic patches of the protein, which are exposed upon thermal denaturation. The thermal denaturation was performed on StepOne Plus Real-Time PCR System (Applied Biosystems). The concentration of mKu was kept constant at 2µl and the DNA concentration was varied from 0 to 8µM. The temperature was ramped from 25° to 95° C with a ramp rate of 1% continuous gradient. The data analysis was performed on the system in-built StepOne Plus software. Tm for the protein was recorded as Boltzmann-derived Tm and Derivative-derived Tm.

### 3.8 Exonuclease assay

The experimental setup was designed based on a previously published protocol to determine the activity of exonucleases [42]. Exonuclease III (Thermo Scientific) was chosen as a standard nuclease for this study at a concentration of 50U/ml. DNA intercalating dye SYBR safe (Invitrogen) was used as the fluorescent reporter to quantify the amount of undigested DNA in each condition. SYBR safe has limited to no fluorescence in solution and it fluoresces only upon intercalating with DNA (λ_ex_/λ_em_ = 502/530nm). The concentration of 40bp dsDNA was kept constant at 0.05µM and mKu dimer was varied from 0 to 0.1µM. The fluorescence signal was detected on Cary Eclipse Fluorescence Spectrometer (Agilent). The excitation filter was set 502nm and emission was detected at 500 to 600nm.

### 3.9 Surface Plasmon Resonance

To assess the impact of DNA length over DNA binding of mKu, surface plasmon resonance experiments were performed in a Biacore T200 SPR machine (Cytiva). For this experiment, Biotin-streptavidin mediated DNA immobilization method was followed [43, 44] on Sensor chip CM-5 (Cytiva). The CM-5 SPR chips are first coated with streptavidin and finally immobilized with biotinylated DNA substrates. The 5’ biotin modified DNA_(65.5%GC)_ substrates of variable lengths 20, 30 and 40 bp were commercially synthesised (Sigma) keeping the sequence unchanged from the *in silico* studies. The supplied biotin tagged single stranded DNA molecules were annealed with an equimolar concentration of the untagged complimentary DNA molecule via heating at 95° C and step-wise cooling to 4°C in a thermal cycler machine. DNA concentration of 1μM was manually injected until ∼200-300 response units (RU) is reached for immobilization to minimize the mass transfer effects. Upon successful immobilization varying concentration of mKu (0-80µM) were flowed until saturation was reached. The concentration dependant responses were processed using the Biacore T200 system inbuilt evaluation software to assess the kinetic properties of the mKu-DNA interaction for each length of DNA substrate. Finally, the data was plotted using OriginPro (Origin Labs) and GraphPad software for further analysis.

### 3.10 Electrophoretic Mobility Shift Assay (EMSA)

EMSA was performed with a commercially synthesised 5’ FAM (Fluorescein amidites) modified DNA (Sigma). For this experiment 40bp DNA_(65.5%GC)_ substrate was used keeping the sequence unchanged from the *in silico* studies (section 3.4). The supplied fluorescent tagged single stranded DNA was annealed with an equimolar concentration of the unlabelled complimentary DNA via heating at 95° C and step-wise cooling to 4°C in a thermal cycler machine. For the DNA binding reaction, 5nM of annealed fluorescent dsDNA product was titrated with 0 to 2.5 and 10nM mKu in a reaction buffer [45] and incubated at 4° C and 25° C for 20 mins and 10mins respectively. The samples were loaded onto a 10% Native PAGE and ran in 0.5x TBE buffer, pH 7.5 at constant 90 V for 2-3 hours. Finally, the gel was visualised using a Fluorescein filter in ChemiDoc MP (Biorad). The imaged gels in triplicates were assessed by densitometric analysis using Imagelab software (Biorad). Data plot, enzyme kinetic and statistical analysis were done on GraphPad Prism software. The generated plot was fitted to non-linear regression model to quantify the equilibrium dissociation constant (K_D_) and hill coefficient (n).

### 3.11 Analytical ultra-centrifugation (AUC)

The oligomeric state of mKu-DNA complex were assessed by determining sedimentation velocity using Optima AUC (Beckman Coulter). DNA, mKu and DNA-mKu samples were spun at 45,000 rpm and monitored in real-time at 288nm wavelength. The raw experimental data was fitted to c(s) model to generate size distribution plot for respective samples.

### 3.12 Negative staining electron microscopy

To visualize the DNA-mKu complexes, 40 bp long dsDNA was commercially synthesised (IDT) keeping the sequence unchanged from the *in silico* studies. DNA and mKu (dimer) was incubated at room temperature for 10-15mins at a equimolar concentration of 15nM. Carbon coated copper grids (PST) were glow discharged at 15mA under vacuum for 30sec. Samples were primarily deposited on the grid for 1 min and blotted dry to remove excess solution. Two subsequent washing step was included with 4ul of nuclease free deionized water followed by staining with 4µl of 2% (v/v) uranyl acetate. Finally, grids were air dried and placed on Technai F30 electron microscope for visualization. A control set with only DNA and only protein were also performed following the same protocol.

## 3. Results

### 3.1 Mycobacterial Ku shares the closest similarity with Omega phage

Multiple sequence alignment was performed with various protein Ku homologues, such as mycobacteriophage omega (ΩKu), human Ku70/80, human Ku70 core DNA binding region (Ku70-cDBR), and few other mycobacterial species (S1 Table 4). mKu shows close sequence similarity with ΩKu (49.1% identity) and Ku70-cDBR (28% identity). Although mKu had limited identity with full-length Ku70/80, due to additional terminal domains (vWA & SAP) of Ku70/80. Moreover, the absence of mKu C-terminal DNA binding stretch (S1 Fig 1) in pathogenic mycobacterial sp. (*M.tuberculosis*, *M.bovis, M.abscessus, M.haemophilium*) was found to be in line with previous reports [17]. Thus apart from other mycobacterial sp., mKu shares the closest sequence similarity with ΩKu.

**Figure 1:**
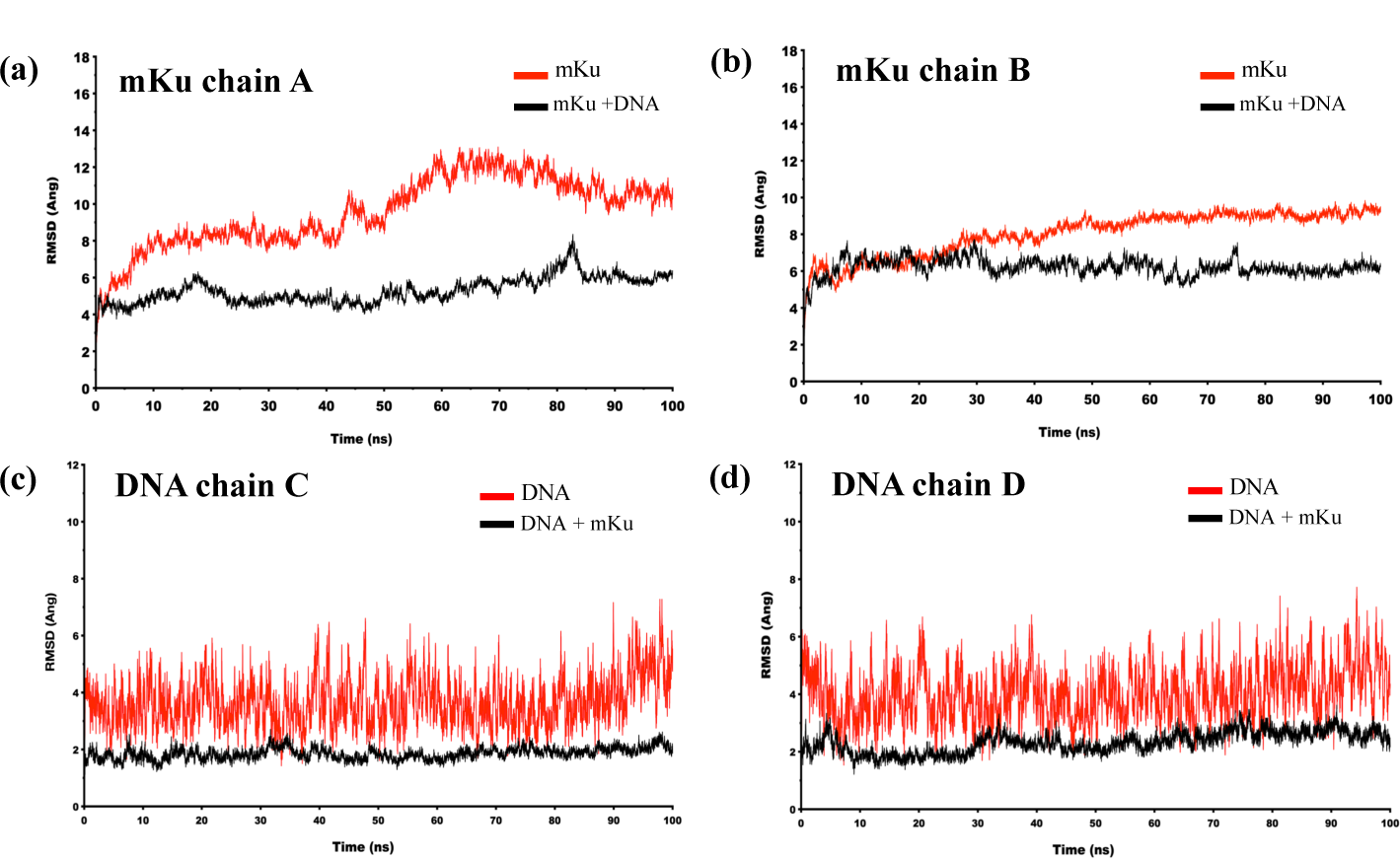
RMSD plots of mKu chains (a & b) and DNA chains (c & d), generated after 100 ns simulation of the modelled structures of mKu dimer, DNA duplex, and mKu-DNA complex. Increased stability of the mKu monomers/chains (a & b) and DNA chains (c & d) is correlated to the decrease in RMSD throughout the simulation.

### 3.2 DNA binding of mKu confers stability to both DNA and the protein

To mimic the genomic composition of *M.tuberculosis*, the GC content of the DNA substrates was kept similar to Mtb genomic DNA at 65.5%. Molecular docking and later MD simulations were performed on mKu-DNA complexes with varying lengths (20, 29 & 40 base pairs). Global stability of each molecule alone and in the complex were assessed by their root mean square deviation (RMSD) throughout 100ns simulation. The average RMSD of mKu chains A and B alone in the dimeric state was 9.5±1.9 Å and 8.3±1.18 Å. While in complex with DNA, a consistent decrease in RMSD was observed for both protein chains, except when in complex with 20bp long DNA. This exception can possibly be due to the total insertion of the 20bp DNA molecule into the inner concavity of mKu dimer with both the DNA ends concealed within the pore. In the case of longer DNAs, mKu was only bound at one end, and the other end of the DNA remained free, as expected physiologically. The mobility of the whole 20bp DNA molecule in the inner concavity may have impacted the stability of the complex(S1.Table 5). To verify this speculation, an hairpin DNA (extracted from PDB: 1JEY) was used as control. Here, the total insertion of the DNA molecule into the mKu dimer was restricted via hairpin formation. In this case, RMSD values of both the protein chains were lowest at 6.33±0.78 Å and 5.74±0.85 Å, when compared with other mKu-DNA complexes. The relative stability of the DNA substrates was parallely assessed in mKu bound and unbound form by their respective RMSDs in a similar fashion (S1.Table 5). In general, the DNA substrates were found to attain additional stability when in complex with mKu, justifying the primary physiological role of mKu to stabilize exposed DNA ends after damage (Fig 1). However, the difference in RMSD for the DNA substrates in apo and complex form seemed to diminish with increasing DNA length (S1.Table 5). From this data, we can predict that a single mKu complex may provide additional stability to DNA molecules (< 40bp). To stabilize longer DNA substrates (>30bp) with two broken ends, more than one mKu dimer may be required, as the movement of the unoccupied broken end might impact the overall stability of the complex.

To study the impact of DNA binding in the structural conformation of mKu. Radii of gyration (Rg) of the backbone residues for the mKu dimer was calculated in apo and DNA bound state (S1.Fig 2) for comparison. The average Rg values for the dimer alone were 26.91 ± 0.48 Å and were increased to 31.20 ± 0.81 Å in the complex form (S1.Table 6). All the mKu-DNA complexes in this study were found to loose compactness upon DNA binding, reflected by their respective increase in Rg (S1.Table 6). From this data, it may be predicted that mKu may adopt two different conformational states in apo and DNA-bound forms, where the apo form has a tighter arrangement. The change in the structural conformation of mKu upon DNA binding may have resulted from additional restraints imposed on the inner concavity of the mKu dimer by the bound DNA double helix. Consistent with the RMSD data, the mKu dimer in complex with 20bp DNA had the highest Rg (31.20±0.81 Å), comparatively more than that of the mKu-DNA_hairpin_ (29.48±0.38 Å), further supporting the impact of 20bp DNA on the stability mKu-DNA complex. Thus, from the MD simulation analysis, it can be predicted that DNA binding of mKu may provide additional stability to both mKu and DNA and may result in loss of compactness of the mKu dimer.

**Figure 2:**
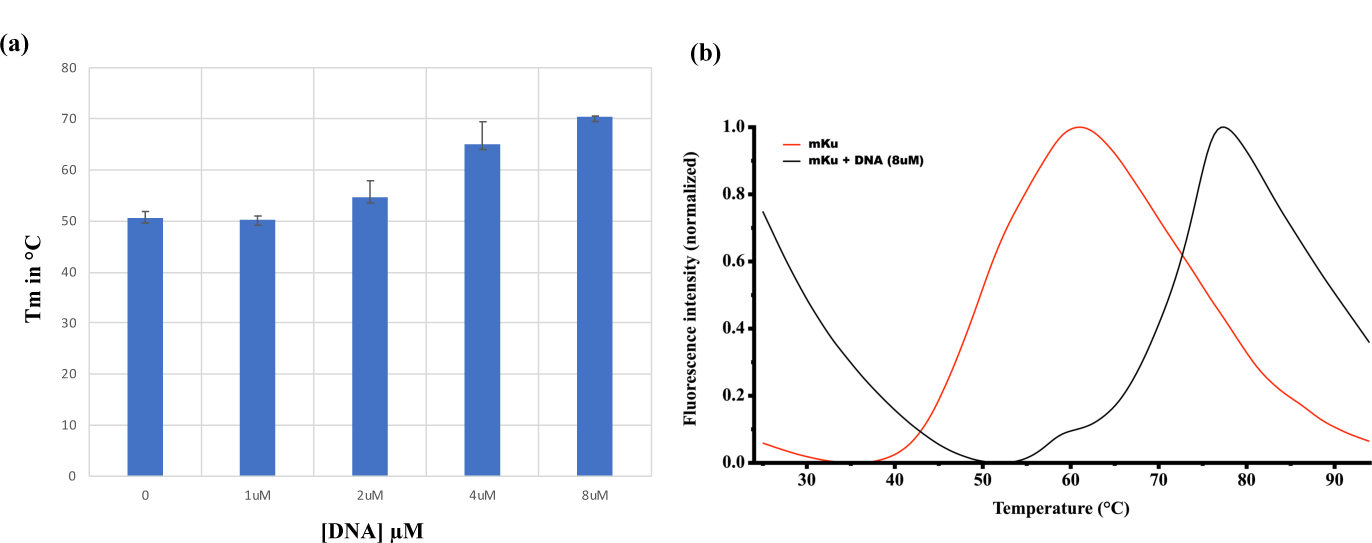
The thermal stability of mKu in the presence and absence of DNA was detected using a solvatochromic fluorescent probe (SYPRO Orange). (a) Mean Tm of mKu graphically represented with increasing DNA concentrations; (b) Melting curve depicting thermal denaturation of mKu with and without DNA.

### 3.3 DNA binding increases the thermal stability of mKu

To validate the impact of DNA binding on the overall stability of mKu, heat denaturation of the protein in the presence and absence of DNA was studied. Using a solvatochromic dye (SYPRO orange) as a reporter, differential scanning fluorimetry was performed on 2µM of mKu dimer with varying concentrations of dsDNA (40bp) from 0 to 8µM. In the absence of DNA, the mean T_m_ (Boltzmann) of mKu was found to be 50.58±1.27 °C (Fig. 2). The high thermal stability of mKu, even in its apo-form, indicates the tight association of the protein dimer. Additionally, from SDS PAGE analysis, we found the mKu dimer to be partially resistant to SDS, which completely denatured upon heating at 100°C for at least 15 mins (S1. Fig 3). The robust nature of the mKu dimer might explain the abundance of the dimeric state of Ku across species in physiological conditions. In the presence of DNA (8µM), the Tm of mKu increased to 70.47±0.11 °C, which correlates to ∼20 °C temperature difference with the apo-form (Fig. 2). Thus, the association with DNA does provide thermal stability to mKu, supporting our previous *in silico* predictions. In humans, Ku70/80 has been reported to get irreversibly entrapped on the DNA, which is later rescued by proteasomes such as p97 [46]. Similarly, mycobacterial Ku might undergo similar irreversible binding with DNA, explaining its resilience to heat and chemical denaturation. Further investigation is required to examine the mKu-DNA interaction and identify whether a similar proteasomal disassembly is involved in post-NHEJ regulation in mycobacteria.

**Figure 3:**
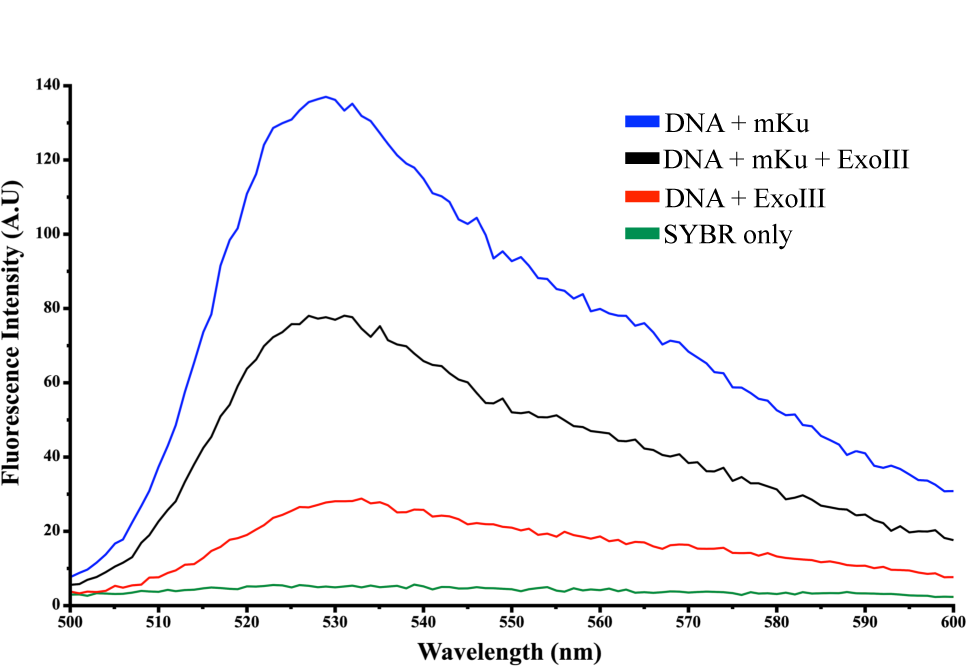
Effect of mKu binding on the nuclease susceptibility of dsDNA. The nuclease activity of ExoIII on 40 bp dsDNA in the presence and absence of mKu was quantified using DNA intercalating fluorescent dye. In the presence of mKu (black), the ExoIII digestion of dsDNA was hindered compared to the reaction mixture with no mKu (red).

### 3.4 Mycobacterial Ku protects the broken DNA ends

DNA ends generated after a double-stranded break are prone to further mechanical and enzymatic damage from Brownian motion and exonucleases present within the cell. Here, we present the potential role of mKu in stabilizing and protecting linear dsDNA. Since the MD simulation study indicated that mKu binding might mitigate the mechanical mobility of the dsDNA. Therefore, we assessed whether the protein can protect dsDNA from enzymatic degradation by an exonuclease assay. The experiment was performed with 0.05µM of 40bp dsDNA and 0.1µM of mKu dimer in the presence and absence of exonuclease III using SYBR safe as the reporter dye. The DNA concentration was kept low to avoid saturation of the fluorescence signal during the assay. SYBR-safe dye alone had minimal to no fluorescence, whereas, in the presence of DNA (negative control), the fluorescence intensity reached around 140 AU (Fig 3). Upon addition of ExoIII (50U/ml), the mixture with no mKu (positive control) showed almost complete digestion of DNA substrate. In the presence of mKu, ∼50% of the signal corresponding to the negative control was retained. It is to be considered that to attain complete protection from ExoIII both the ends of the DNA needs to be capped by mKu, any variation of this arrangement would lead to total or partial nucleolytic digestion. Similarly, in this experiment, the population of DNA molecules with both ends capped by mKu may have attained protection against ExoIII, reflecting the residual fluorescence signal. However, translocation of mKu on DNA will expose the DNA substrate for ExoIII digestion. Thus our data suggests, that mKu might be responsible for DNA stability and end protection against cellular nucleases just after a DSB event. The proposed experimental setup might also help determine the nuclease sensitivity of other end-binding DNA-protein complexes.

### 3.5 DNA length impacts the binding kinetics of mKu-DNA

*In silico* studies on the mKu-DNA complex via MD simulation suggested that the overall stability of the complex might be affected by the length of the DNA (S1 Table 5). To explore this, we looked into the DNA binding kinetics of mKu with different lengths of DNA (20, 29, and 40 bp), keeping the nucleotide sequence unchanged from the *in silico* experiments. The kinetic experiments were performed with the help of surface plasmon resonance (SPR) using immobilized biotinylated dsDNAs and subjecting it to varying concentrations of mKu (0-80µM). From the sensograms corresponding to the respective DNAs (Fig 4.b - d), it was found that the magnitude of response (RU) generated from mKu binding is dependent on the DNA length. The Vmax for 20, 29 and 40 bp DNA substrates were 43.39±3.23, 261.9±25.17, and 863.1±206.6 RU, respectively, suggesting higher occupancy mKu molecules on longer DNA substrates. Similarly, to determine the binding characteristics of the interaction, the plots were transformed as a function of mKu concentration and fitted to the Michaelis-Menten kinetics model (S1 Fig 4).

**Figure 4:**
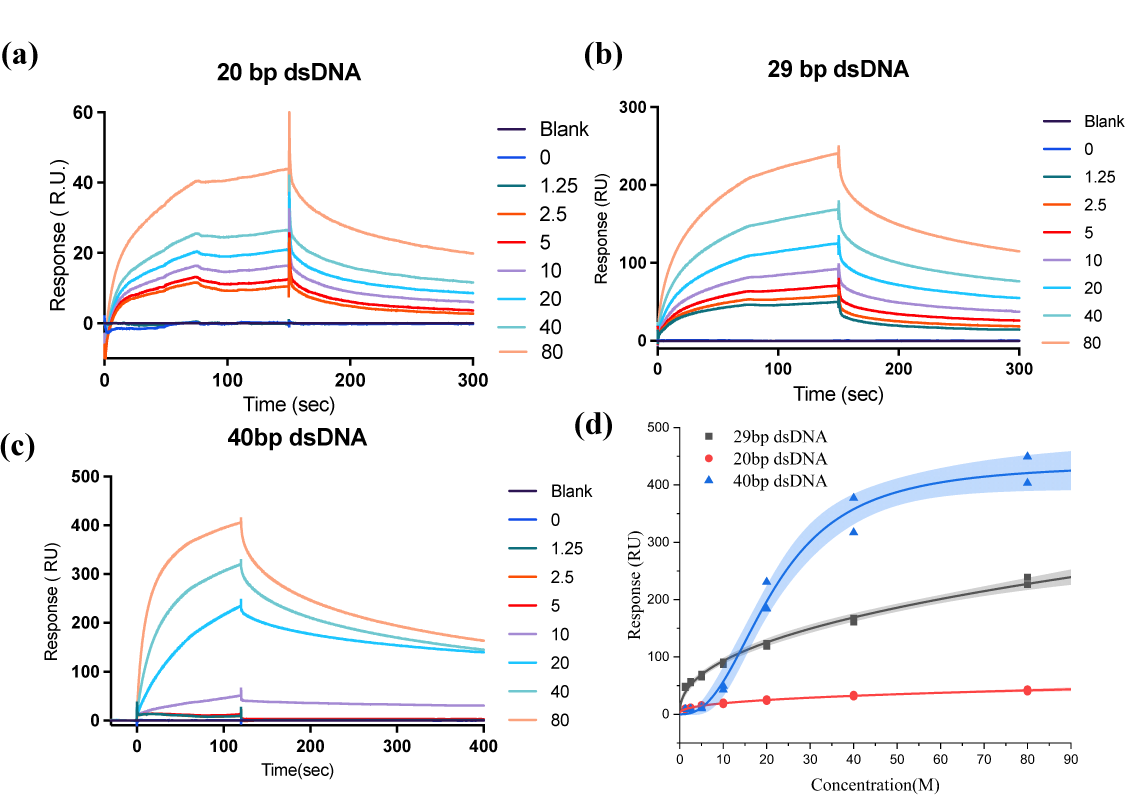
DNA length-dependent binding kinetics of mKu: SPR sensograms for 20bp(a), 29bp(b), and 40bp(c) long DNA substrates with varying concentrations of mKu (0-80nM). Legend provided on the right (a, b & c) denotes mKu concentrations for each curve in nM; Concentration dependent kinetics of mKu (d) on multiple (20, 29 & 40 bp) DNA substrates fitted to Hill model with R^2^ of 97.5, 98.8 and 98.5%, respectively.

The calculated K_M_ for 20, 29, and 40bp DNA were 10.79±2.47, 17.86±4.58, and 74.34±30.25 nM, respectively. A single Ku (human) molecule is reported to span an equivalent area of 18 to 20 bp long DNA [47]; thus, a 40bp long DNA may accommodate at least two or more mKu molecules. In this experimental setup, one end of the DNA was tethered to the SPR chip, and it is inaccessible to mKu for binding. Thus, we speculate that, following the initial binding of one mKu dimer at the free DNA termini, the bound mKu molecule must slide on the DNA for the entry of the next molecule from the accessible end of the DNA strand.

We further investigated the cooperativity of mKu-DNA interaction by fitting the binding kinetics to the Hill model. The obtained Hill coefficient (n) for DNA substrates were 0.38±0.09 (20bp), 0.43±0.07 (29bp), and 2.45±0.34 (40bp), where n=0 indicates no cooperativity, n < 1 indicates negative cooperativity and n > 1 indicates positive cooperativity. Therefore, it can be inferred that the length of the DNA does impact the DNA-protein interaction, and mycobacterial Ku has positive cooperativity for dsDNA of length ≥ 40bp; this finding is in line with the human Ku70/80 [47]. Furthermore, mKu might be able to translocate on the DNA long enough to accommodate more than one mKu molecule. It has to be acknowledged that in this SPR experimental setup, one end of the DNA was inaccessible to mKu due to immobilization on the SPR chip. Therefore, the binding affinity obtained from this experiment may not reflect the natural binding affinity of DNA end-binding proteins. Since Ku is known to bind specifically to DNA ends thus, the calculated K_D_ via SPR might be higher than the actual value. Complimentary methods, such as EMSA, may better quantify the binding affinity. Nonetheless, to comparatively study the impact of DNA length on the binding kinetics of DNA-protein interaction, the SPR-based experimental setup has been considered reliable and extensively used [43, 44].

Electrophoretic mobility shift assay (EMSA) was performed to investigate the DNA-mKu interaction. The assay was performed with a fixed concentration of fluorescently labeled 40 bp long DNA titrated with 0 to 2.5 nM (Fig 5.a) and 10 nM (S1.Fig 5) mKu to quantify the DNA binding affinity of the protein. Relative intensities of the bands on the gel were determined from densitometric analysis. The total DNA (Fig 5.a) without any protein was considered as the total unbound DNA (U_T_). Titration of the DNA with increasing concentration of mKu, resulted in a gradual decrease of the unbound DNA fraction (U_x_), suggesting the formation mKu-DNA complex, which can be visualized at the top section of the lane (Fig 5.a). The difference in the gel migration of the DNA and the complex corresponds to their respective molecular weight. The band intensities of U_T_ and U_x_ were used to calculate and plot the relative fraction of bound DNA as a function of mKu concentration (Fig 5.b). The generated plot was fitted to a non-linear regression model, and the Hill coefficient (n) was calculated to be 1.5±0.3, suggesting positive cooperativity. The equilibrium dissociation constant (K_D_), denoting the concentration of mKu where 50% of the total DNA is bound, was found to be 0.33 ± 0.08 nM with B_Max_ of 1.516 ± 0.414 nM in a 95% likelihood profile. The DNA binding affinity of mKu is comparable to earlier reports on Human Ku70/80 (K_D_ = 0.38 nM) [47]. The strong affinity of Ku for linear DNA justifies the earlier reports on the entrapment of the Ku on DNA [46].

**Figure 5:**
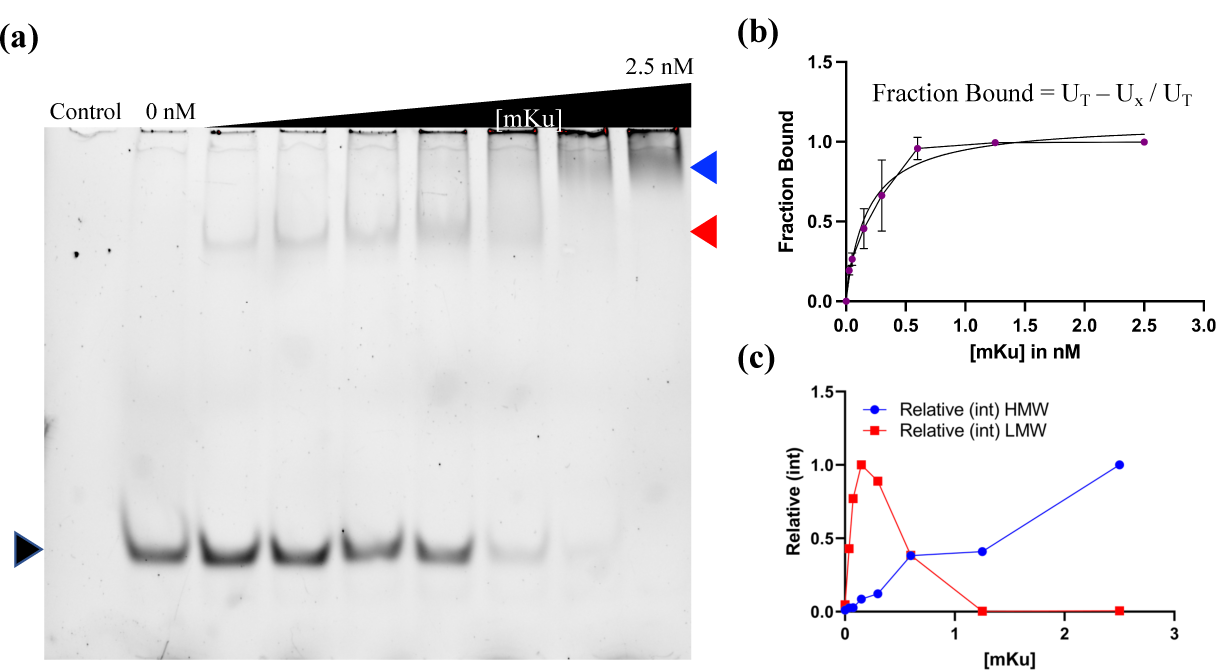
Gel shift assay of mKu (0 to 2.5nM) binding to 5’FAM tagged dsDNA (5nM). (a) Gel image showing free DNA (black arrow) and mKu-DNA complexes of different sizes (red and blue arrow); (b) Graphical representation of the bound DNA with increasing concentration of mKu, calculated from the band intensities (black arrow); (c) Relative intensities of high and low molecular weight mKu-DNA complexes (LMW & HMW).

Interestingly, the DNA-protein complex of two different sizes (indicated with red and blue arrows) can be spotted on the gel (Fig 5). For simplicity, we termed the complexes, as per their relative molecular size, as low and high molecular weight complexes (LMW & HMW). The relative disappearance and emergence of LMW and HMW were found to be dependent on the concentration of mKu (Fig 5.c). Multiple oligomeric states of the mKu-DNA complex can be speculated to be caused by the binding of more than one mKu to the DNA with increasing concertation of mKu, supporting our findings from SPR experiments.

### 3.6. mKu-DNA complexes form higher oligomeric assemblies

To estimate the molecular size of the DNA-mKu complexes, we have performed size exclusion chromatography (SEC) on Superdex 200 Increase 10/300 GL. The ratio of mKu dimer to 40 bp DNA concentration were kept at 2:1. The protein alone was eluted at ∼20ml whereas the mKu-DNA complex was eluted at 9 ml and 15 ml. Complexes with two different sizes as seen in the SEC chromatogram (Fig 7.b), is in agreement with the gel-based assay (Fig 5.a). To further support and quantify the size of the complex, analytical ultra-centrifuge was performed on mKu, DNA, and mKu-DNA complex.

**Figure 6:**
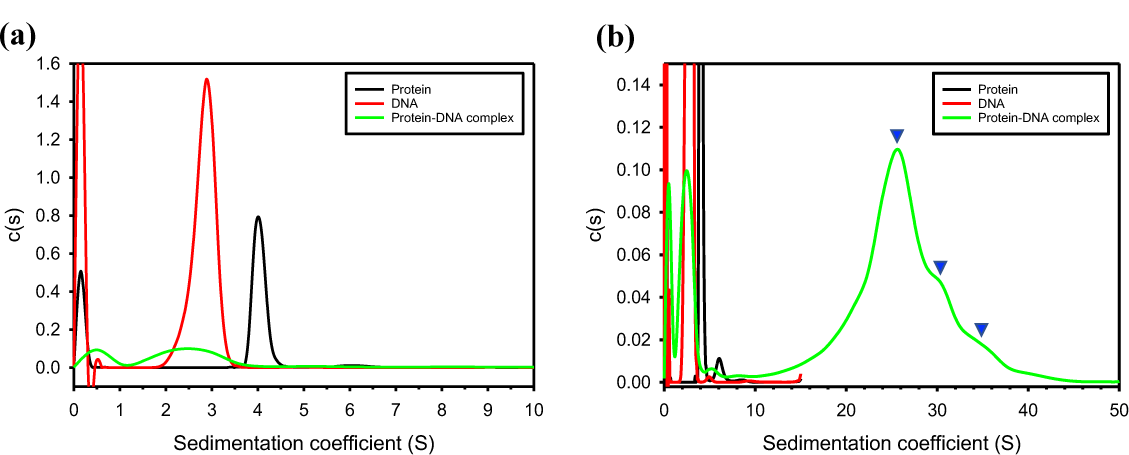
Size distribution of mKu protein (black), DNA (red), and mKu-DNA complex (green) shown as a function of sedimentation coefficient (S), obtained using analytical ultra-centrifugation. (a) and (b) represent a single size distribution plot focused at two windows along X-axis for better visualization. mKu-DNA complex may have higher order oligomeric states upon association of each unit complex, as shown with blue arrows in (b).

**Figure 7:**
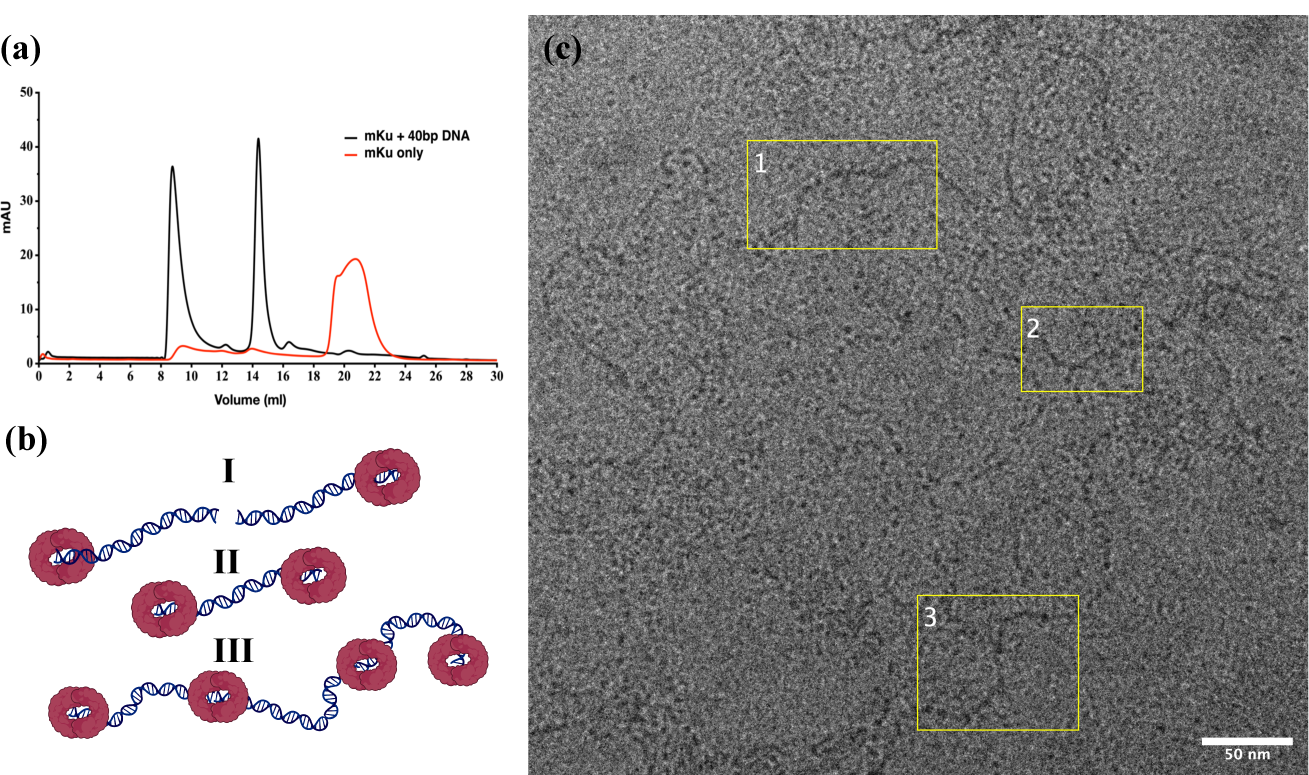
Higher order oligomerization of mKu-DNA (40 bp) complex: SEC profile representing elution profile of mKu only and mKu + 40bp DNA (a); Schematic showing possible oligomeric assemblies of mKu-DNA complex leading to the formation of filament-like structures (b). mKu homodimer (shown in maroon) is shown to assemble on DNA in one-to-one (b.I), two to one(b.II), or end-to-end assembly (b.III) by interacting with two DNA ends; mKu bound DNA oligomers as seen positively stained with uranyl acetate, under TEM at 78KX magnification. Oligomeric assemblies in the image are emphasized (yellow boxes) as linear (c.1&3) and branched structures (c.2)

The sedimentation coefficient for protein, DNA, and the complex obtained from the size distribution plot was found to be 4 S, 2.8 S, and 25.3 S, respectively (Fig 6). The coefficient for mKu and DNA alone aligned with their theoretical sizes, but it was unexpectedly higher for the complex. Considering the 2:1 stoichiometry of mKu dimer to DNA, the complex’s expected sedimentation coefficient should be less than 10S, unlike the experimental value of ∼25S. As the 40 bp long DNA cannot physically accommodate more than two to three mKu molecules, it can be speculated that the individual unit complexes may have interacted to form higher oligomers, as depicted in the schematic representation (Fig 7.b). Moreover, the peak for the mKu-DNA complex (Fig 6.b) was found to be heterogenous, indicated by the shoulder peaks. To confirm that the oligomeric assembly is an ordered association, unlike randomly formed aggregates, we visualized the sample under the transmission electron microscope (TEM). Linear long filament-like structures were seen with negative staining of the mKu-DNA complex. The length of these filaments was orders of magnitude higher than the single 40 bp DNA stands (12.56 nm). Thus, we predict two possibilities which may explain long filament-like oligomers of mKu-DNA complex. First scenario where each dumble-like mKu-DNA complex with 2:1 stoichiometry participate in an end-end association to form a long filament-like assembly, primarily governed by cross mKu dimer interaction (Fig 7.c). Although no cross dimer-dimer interaction or filament-like structures were seen for mKu in absence of DNA. Secondly, due to the symmetric nature of mKu dimer, DNA can ideally lodge from both ends of the DNA binging pore. In this case, one protein dimer may bind to two dsDNAs from the opposite ends of the DNA binding pore. The ability of mKu to form higher oligomers by interacting with two dsDNAs may shed some light on its role in mycobacterial NHEJ synapsis, similar to previous reports on *B.subtilis* [48]. DNA synapse is a critical step in DSBR, as initiating the end-joining process depends on the spatial distance between broken ends, which is otherwise under random motion due to diffusion. The mechanism of NHEJ synapsis in prokaryotes is unclear due to the absence of proteins like XLF, which are reported to actively participate in human NHEJ synapsis along with Ku and Ligase IV [49, 50]. Although the end-to-end association of the mKu-DNA complex requires further scrutiny, our current data suggest that mKu and ligase D might be responsible for bringing broken ends of DNA together, forming mycobacterial NHEJ synapsis.

## 4. Discussion

NHEJ is the sole double-stranded break repair machinery for *Mycobacterium tuberculosis* during dormancy, aiding its survival and sustenance in the host macrophage. Mycobacterial NHEJ pathway is a two component system comprising of Ku homodimer and an ATP dependant ligase (Ligase D). DNA end binding protein Ku is considered to be the rate limiting protein due to its ability to simultaneously bind to the broken dsDNA and to ligase D forming the NHEJ complex. Thus, we have focused on mycobacterial Ku (mKu) to better understand DSBR in *M.tuberculosis*. In this study, we have exploited *in silico* and *in vitro* tools to understand the DNA binding properties and delineate the potential roles of mKu in mycobacterial NHEJ. From molecular dynamics simulation of mKu homodimer model with and without DNA, we found that the DNA binding of mKu imparts stability to both DNA and mKu dimer (Fig 1). To validate our *in silico* finding, we have employed differential scanning fluorimetry (DSF) to assess the thermal stability of mKu in presence and absence of DNA. The difference in melting temperature for mKu homodimer alone and with dsDNA (40bp) was found to be ∼20°C (Fig 2), supporting our *in silico* data. Further, we examined whether binding of mKu can protect the broken DNA ends from exonuclease digestion, which are otherwise prone to degradation by cellular nucleases. To assess this, we have designed and performed a fluorescence based nuclease assay with mKu and DNA in presence of fluorescent DNA intercalator. Interestingly, we found that ∼50% of the total DNA to be protected in presence of mKu. Although the concentration mKu used for the assay was twofold higher as compared to DNA, which in theory should confer complete resistance to the bound DNA from exonuclease digestion. The partial sensitivity of mKu bound DNA may be explained by mKu’s ability to slide on the DNA, which might expose the free DNA ends to ExoIII. DNA sliding phenomena of human Ku70/80 has been previously reported [47] and we have established the same for mKu from the SPR based experiment (section 4.5). To assess whether DNA length can impact the DNA binding kinetics of mKu, we have performed SPR on mKu and dsDNA with various lengths (20, 29 and 40bp). mKu showed the highest affinity for the shortest DNA (20bp) corresponding to the lowest K_M_ among all the DNA substrates (Fig 4). Additionally, mKu showed positive cooperativity for dsDNA ≥40bp in length, which is in agreement with human Ku70/80 [47]. From R_max_ corresponding to individual DNA substrates (Fig 4.d), we found the longest DNA (40bp) to have the highest occupancy of mKu. We speculate that the variable occupancy of mKu molecules on DNA depending on the DNA length is the result of mKu’s translocation on the DNA following end binding. Thus, our SPR based study suggests the sliding of mKu dimer on DNA, similar to human Ku70/80 [47].

To quantify the DNA binding affinity and assess the homogeneity of the mKu-DNA complex, we have performed electrophoretic mobility shift assay (EMSA). mKu showed strong DNA binding affinity for 40bp long DNA with equilibrium dissociation constant (K_D_) of 0.33±0.08 nM, which is close in comparison to human Ku70/80 [47]. Additionally, we have observed mKu-DNA complex to exist in multiple sizes in gel (Fig 5.a & c). Thus, to closely determine the oligomeric states of mKu-DNA complex we performed analytical ultra-centrifuge(AUC). The mKu-DNA complex was found to be heterogenous with more than one oligomeric state supporting the our observations from EMSA (Fig 6). The sedimentation coefficient for the complex was calculated to ∼25S, which was unexpectedly higher than theoretical value of mKu-DNA complex in 2:1 stoichiometry. Therefore, we speculate that mKu may form higher order oligomers by interacting with more than one molecule of DNA per mKu dimer. The symmetric nature of the DNA binding pore of mKu dimer at two opposite ends may allow mKu to associate with two DNA molecules (Fig 7.a). To support our speculation we visualized the mKu-DNA (40bp) complex under transmission electron microscopy. The protein was seen to form long filament-like structures with DNA, which are orders of magnitude larger the individual DNA substrate (Fig 7.c). The ability of mKu to form continuous filament like structures with DNA may indicate its role in bringing broken DNA ends in proximity, an event also known as NHEJ synapsis. Mycobacterial NHEJ lacks the accessory factors which are devoted solely to DNA synapsis in human[51]. Although the role of mKu in NHEJ synapsis is subject to further examination but our current data suggests that mKu can actively bring free DNA ends in close proximity, forming filament-like higher order oligomeric structures.

## 5. Conclusion

Our study focuses on the DNA end binding protein (mKu) of mycobacterial NHEJ and closely inspects its interaction with DNA and its possible role in DNA synapsis, using *in silico and in vitro* tools. The molecular dynamics simulation, complemented with *in vitro* experiments, suggests that the association of mKu with DNA (at the termini) imparts additional stability to both mKu and DNA, when compared to their respective apo/unbound form. From the R_g_ (radius of gyration) of the complexes obtained from MD simulation, mKu was predicted to exist in two conformational states (compact and loose) in bound and unbound form, respectively. EMSA and SPR studies suggest that mKu has a high affinity for linear dsDNA and shows positive cooperativity with longer DNA substrates(≥40bp). mKu was found to form a higher-order oligomeric assembly with DNA(≥40bp). We speculate that the oligomerisation may be the result of binding more than one DNA molecule per mKu dimer due to the symmetric nature of its DNA binding pore. The ability of mycobacterial Ku to lodge two DNA molecules at opposing ends indicates its potential role in bringing two free DNA ends together, forming synapsis. However, mycobacterial NHEJ needs to be studied in more detail to support the potential role of mKu in DNA synapsis.

### Authorship contribution statement

JB conceptualized, designed and performed the experiments. JB performed the downstream data analysis and wrote the manuscript. GB performed the MD simulation. SD performed the SPR assay. DS supervised SD and JB in SPR data analysis. AKD, IR and EH helped in conceptualization and supervised the research. All authors have read and approved the manuscript.

## Supporting information

https://docs.google.com/document/d/10F-GO27lC4B4Ikof7Wy2AZD1utk_HMnr/edit?usp=share_link&ouid=116619428028898377979&rtpof=true&sd=true

## Acknowledgements

We thank the Melbourne India postgraduate academy (MIPA) program for the joint collaboration between IIT Kharagpur and university of Melbourne. This work was financially supported by the core research grant (DST No. CRG/2020/002622) granted to Amit Kumar Das, sanctioned by science and engineering research board (SERB), Government of India. National Health and Medical Council (NHMRC) Ideas APP2000934 and the start-up package of the University of Melbourne granted to Isabelle Rouiller. We thank Dr. Yee-foong Mok, Melbourne protein characterization facility, University of Melbourne for his help with AUC experiments. We also thank Prof. Aidan Doherty, University of Sussex, for the gift of the recombinant plasmid containing mKu (pET16b-mKu) used in this study.

