## Supplementary material for "*In silico* and *in vitro* characterization of the mycobacterial protein Ku to unravel its role in non-homologous end-joining DNA repair": https://docs.google.com/document/d/10F-GO27lC4B4Ikof7Wy2AZD1utk_HMnr/edit?usp=share_link&ouid=116619428028898377979&rtpof=true&sd=true

***Supplementary: Tables***

**S1 Table 1:** Comparative overview of predicted models from multiple web servers

| Web server | RMSD from Ku70-cDBR | Allowed residues | Favoured residues | Ramachandran outliers |
|---|---|---|---|---|
| Ku70-cDBR (PDB: 1JEY) | 0 | 98.2% | 86.8% | 5 |
| Swiss-Model | 0.938 | 99.2% | 89.7% | 2 |
| Robetta | 3.754 | 99.3% | 97% | 2 |
| Phyre2 | 1.839 | 97.8% | 92.3% | 6 |
| I-Tasser | 1.161 | 93.4% | 78.6% | 18 |

**S1 Table 2:** Overview of predicted mKu homodimer from various molecular docking servers

| Web Server | Van der Waals potential(kcal/mol) | Solvation free energy(kcal/mol) | Total Binding free energy (kcal/mol) |
|---|---|---|---|
| Cluspro | -306.79 | -42.31 | -193.05 |
| Haddock | -119.45 | -17.41 | -26.08 |
| ZDock | -186.52 | -25.95 | -93.06 |

**S1 Table 3:** Details of all the DNA substrates used in this study

| **Identifier** | **Length (bp)** | **Properties** | **Sequence** 5'→ 3' | **Origin** | **%GC content** |
|---|---|---|---|---|---|
| **DNA$_{1JEY}$** | 21 & 34 | Double helix with 3' hairpin loop | GCCCAGCTTTCCCAGCTAATAAACTAAAAAC<br><br>GTTTTTAGTTTATTGGGC | Extracted from Ku70/80 hetero-dimer DNA complex (PDB:1JEY) | 33 & 41 |
| **DNA$_{(20bp)}$** | 20 | Double helix | GTCGTAGCCCACCTGACCGT<br><br>ACGGTCAGGTGGGCTACGAC | Random generated | 65.5 |
| **DNA$_{(29bp)}$** | 29 | Double helix | CGAACCAGCTGGCGCATAGCCGCGCTATC<br><br>GATAGCGCGGCTATGCGCCAGCTGGTTCG | Random generated | 65.5 |
| **DNA$_{(40bp)}$** | 40 | Double helix | CCCCCCTGTCGCCGCCGACGTCTGTGATATGGCGTTGTTG<br><br>CCCCCCTGTCGCCGCCGACGTCTGTGATATGGCGTTGTTG | Random generated | 65.5 |

**S1 Table 4:** Multiple sequence alignment of mKu with homologues from other species

| Input Sequence | Coverage | Percentage Identity |
|---|---|---|
| mKu and Omega phage Ku | 91.1% | 49.1 % |
| mKu and Human Ku70/80 | 74.5 and 32.2% | 15.1% and 8% |
| mKu and Human core DNA binding region (Ku70) | 95% | 28% |
| mKu , Ku (*M.bovis*) and Ku (*M.Smegmatis*) | 100% | 73.3 and 64.6% |

**S1 Table 5:** Average RMSD (Å) over 100ns of MD simulation for multiple DNA-mKu complexes

| Complex | mKu Chain A | | mKu Chain B | | DNA Chain C | | DNA Chain D | |
|---|---|---|---|---|---|---|---|---|
| | Unbound | Bound | Unbound | Bound | Unbound | Bound | Unbound | Bound |
| mKu+1JEY | 9.58±1.9 | 6.33±0.78 | 8.03±1.18 | 5.74±0.85 | 3.65±0.82 | 1.85±0.20 | 4.0±0.92 | 2.32±0.37 |
| mKu + 20 bp | | 10.38±1.91 | | 8.62±1.5 | 4.85±1.01 | 2.34±0.45 | 5.29±1.11 | 3.25±0.95 |
| mKu + 29 bp | | 7.27±0.78 | | 8.08±0.80 | 6.07±1.13 | 4.95±0.92 | 6.24±1.39 | 4.95±1.05 |
| mKu + 40 bp | | 5.30±0.70 | | 6.21±0.49 | 7.27±1.08 | 8.50±1.53 | 6.72±1.03 | 7.39±1.47 |

**S1 Table 6:** Average radius of gyration of mKu and mKu-DNA complexes

| Complex | mKu dimer (Unbound) in Å | mKu-DNA complex (Bound) in Å |
|---|---|---|
| 1JEY | 26±0.48 | 29.48±0.38 |
| 20 | | 31.20±0.81 |
| 29 | | 27.51±0.53 |
| 40 | | 28.05±0.38 |

### *Supplementary: Figures*

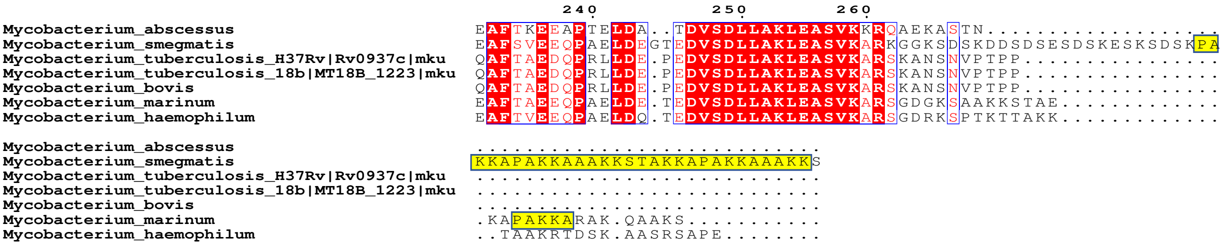


**S1 Figure 1**: Multiple sequence alignment of mKu from *M.tuberculosis* with pathogenic and non-pathogenic variants of Mycobacteria sp.: C-terminal 'PAKKA' DNA binding repeat sequence (highlighted in yellow) can be seen to be absent in pathogenic variants of mycobacteria such as, *M.tuberculosis*, *M.bovis*, etc.

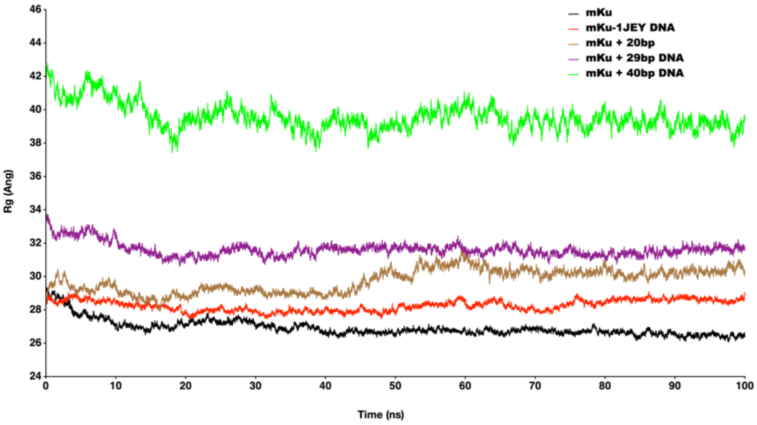


**S1 Figure 2**: Radius of gyration of mKu and mKu-DNA complexes representing overall compactness of each complex over 100ns of MD simulation

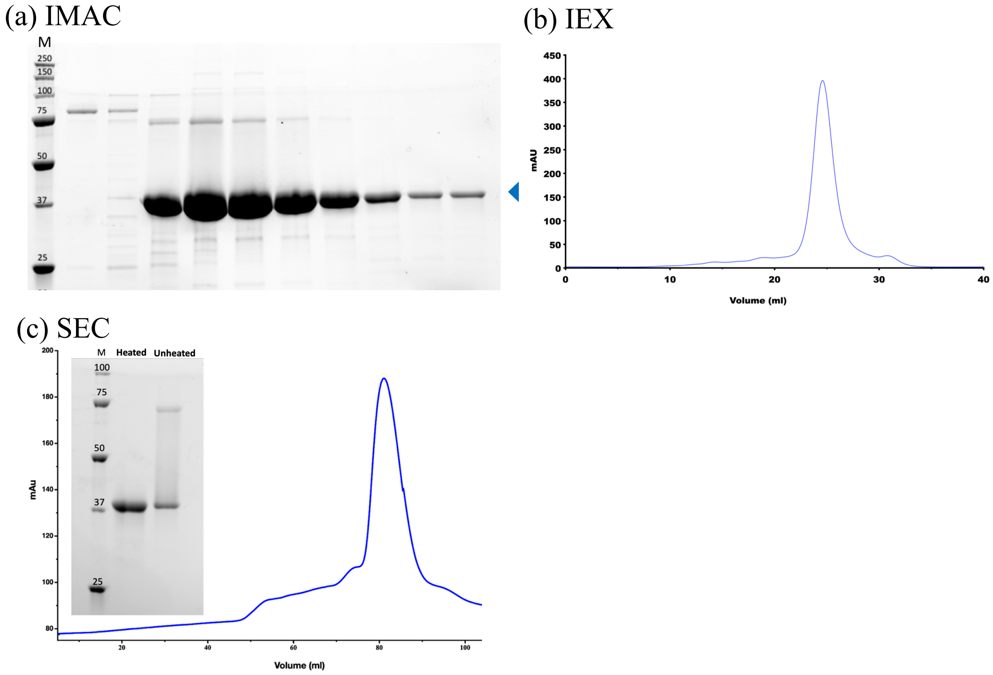


**S1 Figure 3**: SDS PAGE of elution fractions of mKu (left to right) from Ni-NTA chromatography(a), the band corresponding to mKu is marked with blue arrow; Anion exchange chromatograph of Ni-NTA eluted fractions(b); Size exclusion chromatograph of mKu(c) showing elution of mKu at ~40ml; SDS PAGE showing partial SDS resistance mKu dimer in heated and unheated condition(d).

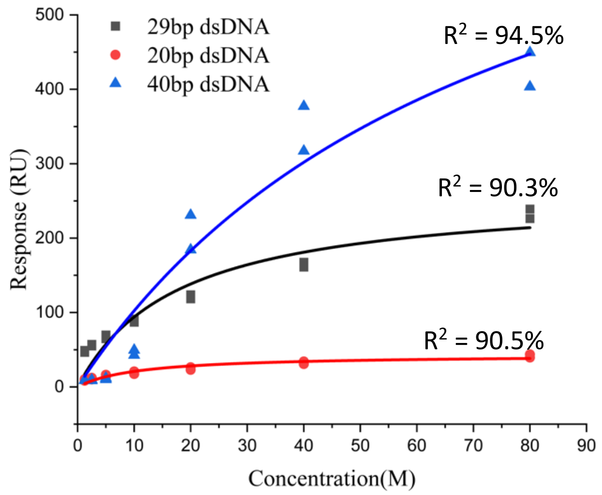


**S1 Figure 4**: Concentration dependent kinetics of mKu on 20bp, 29bp & 40bp DNA substrates fitted to Michaelis-Menten kinetics model (95% confidence).

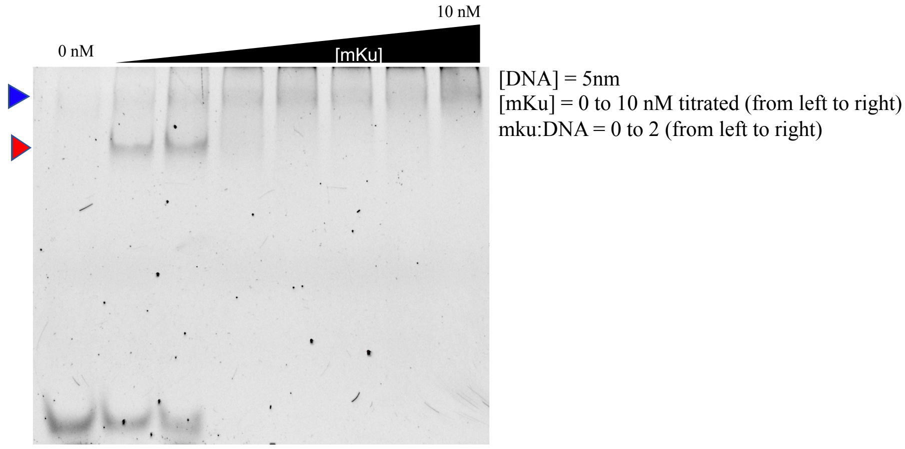


**S1 Figure 5**: EMSA of mKu(0-10nM) binding to 40bp DNA showing unbound DNA fractions (bottom), DNA-mKu complexes of different sizes shown with blue and purple arrows.
